# Thresholding Approach For Low-Rank Correlation Matrix Based On Mm Algorithm

**DOI:** 10.1101/2021.12.28.474401

**Authors:** Kensuke Tanioka, Yuki Furotani, Satoru Hiwa

## Abstract

**Background:** Low-rank approximation is used for interpreting the features of a correlation matrix using visualization tools; however, a low-rank approximation may result in estimation that is far from zero even if the corresponding original value is zero. In such a case, the results lead to a misinterpretation.

**Methods:** To overcome this, we propose a novel approach to estimate a sparse low-rank correlation matrix based on threshold values. We introduce a new cross-validation function to tune the corresponding threshold values. To calculate the value of a function, the MM algorithm is used to estimate the sparse low-rank correlation matrix, and a grid search was performed to select the threshold values.

**Results:** Through numerical simulation, we found that the false positive rate (FPR) and average relative error of the proposed method were superior to those of the tandem approach. For the application of microarray gene expression, the FPRs of the proposed approach with *d* = 2, 3, and 5 were 0.128, 0.139, and 0.197, respectively, whereas the FPR of the tandem approach was 0.285.

**Conclusions:** We propose a novel approach to estimate sparse low-rank correlation matrices. The advantage of the proposed method is that it provides results that are interpretable through the use of a heatmap, thereby avoiding result misinterpretations. We demonstrated the superiority of the proposed method through both numerical simulations and real examples.

## 1 Background

When describing the linear relationship between two variables or subjects, a correlation matrix is calculated from multivariate data. For example, in the domain of genomics, a correlation matrix between genes is used in combination with a heatmap [Wilkinson and Friendly, 2009]. However, network structures of these correlation coefficients are masked because real data contain noise.

To address this problem, a low-rank approximation is used[ten Berge, 1993]. For the estimation of low-rank correlation matrices, various methods have been proposed [Pietersz and Groenen, 2004, Simon and Abell, 2010, Grubisic and Pietersz, 2007, Duan et al., 2016], and there are two advantages associated with this estimation. First, it becomes easy to interpret the network structure of the correlation matrix visually because variations in low-rank correlation coefficients tend to be larger. Therefore, the heatmap of the estimated low-rank correlation matrix provides a readable visualization. Second, low-rank approximations also can effectively describe the clustering structure [Ding and He, 2004], which results in improved interpretations. Finally, it must be noted that the values of the estimated low-rank correlation coefficient can range from 1 to 1, in contrast to that of singular valued decomposition (SVD). Therefore, the relationships between the variables can be easily interpreted using low-rank approximations because the obtained values are bounded.

It is indeed the case that low-rank correlation matrices simplify relations; however, there are several problems with such approximations. Because of the advancements in the field of information technology, the amount of data available is quite large. Therefore, the number of coefficients needed for interpretation from such large amounts of data is above the cognitive boundary. Second, even if a true correlation coefficient is close to zero, the corresponding estimated low-rank correlation coefficient can be far from zero. The estimated results will lead to a misinterpretation of the relationships between the variables.

To overcome this, in this study, we proposed a new approach to estimate sparse low-rank correlation matrices. The proposed approach, combined with a heatmap, provides a visual interpretation of the relationships between the variables. For the sparse methods of the correlation matrix and covariance matrix [Engel et al., 2017, Lam, 2020], there are two types of methods that are available. The first involves adding a sparsity penalty to the objective functions [Bien and Tibshirani, 2011, Rothman, 2012, Xue et al., 2012, Cai et al., 2011, D’aspremont et al., 2008, Friedman et al., 2008, Rothman et al., 2008, Yuan and Lin, 2007, Cui et al., 2016]. The other type uses thresholding values to achieve a sparse structure. Bickel and Levina [2008a] proposed the thresholding matrix estimator and various related methods have been developed [Cai and Liu, 2011, Bickel and Levina, 2008b, El Karoui, 2008, Jiang, 2013]. In addition, to estimate sparse correlation matrix, [Rothman et al., 2009, Lam and Fan, 2009, Liu et al., 2014] used generalized thresholding operator based methods [Rothman et al., 2009], respectively. For the estimation of sparse low-rank matrices, methods based on penalty terms have also been proposed [Zhou et al., 2015, Savalle et al., 2012].

The proposed approach adopts an approach that uses hard thresholding based on Bickel and Levina [2008a] and Jiang [2013] therefore has the ease of use and provides interpretable results Therefore, to estimate sparse low-rank correlation matrices, we introduce a new cross validation function, a modification of those used in Bickel and Levina [2008a] and Jiang [2013]. To calculate the values of the cross validation function, the majorize-minimization algorithm (MM algorithm) [Pietersz and Groenen, 2004, Simon and Abell, 2010] and the hard thresholding approach are used. The proposed approach has two advantages; the estimated sparse low-rank correlation matrix allows for an easy and visual interpretation of the correlation matrix using a heatmap and avoids the misinterpretation of the correlation matrix. When the true correlation coefficient is zero, the proposed method tends to estimate the corresponding coefficient as zero. In addition, we focus only on positive correlation coefficients, not negative correlation coefficients. With the focus on only positive relations, it becomes easy to interpret the features of relations.

The rest of this paper is structured as follows. We explain the model and algorithm in section 2. In section3, we evaluate the proposed approach and describe the numerical simulation. The results of applying the proposed method to real data are provided in Section 4. We finally present our conclusions in Section 5.

## 2 Method

### 2.1 Adaptive thresholding for sparse and low-rank correlation matrix estimation

In this section, we present the proposed approach for estimating a sparse low-rank correlation matrix. First, an estimation of a low-rank correlation matrix is described based on the MM algorithm [Pietersz and Groenen, 2004, Simon and Abell, 2010]. Next, to achieve the sparse low-rank correlation structure, the hard thresholding operator and proposed cross-validation function are described.

#### 2.1.1 Optimization problem of low-rank correlation matrices

Let ***R*** = (*r*_*ij*_) *r*_*ij*_ ∈ [−1, 1] (*i, j* = 1, 2, · · ·, *p*) and ***W*** = (*w*_*ij*_) *w*_*ij*_ ∈ {0, 1} (*i, j* = 1, 2, · · ·, *p*) be correlation matrix between variables and the binary matrix, respectively, where *p* is the number of variables. Given the number of low dimensions *d* ≤ *p* and correlation matrix ***R***, binary matrix ***W***, the optimization problem of estimating a low-rank correlation matrix is defined as follows.

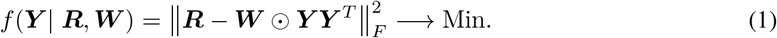

subject to

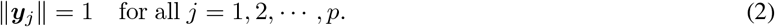

where ***Y*** = (***y***_1_, ***y***_2_, · · ·, ***y***_*p*_)^*T*^, ***y***_*j*_ = (*y*_*j*1_, *y*_*j*2_, · · ·, *y*_*jd*_)^*T*^, 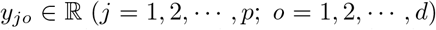 is coordinate matrix of variables on *d* dimensions, ⊙ is the Hadamard product, and ∥ · ∥_*F*_ is Frobenius norm. The objective function in Eq. (1) was explained by Knol and Ten Berge [2012]. From the constraint (2), ***Y***^*T*^***Y*** becomes the correlation matrix. When ***Y*** is estimated, ***W*** is fixed. In the situation, ***Y*** is estimated based on MM algorithm. The estimation is described in 2.1.2. For the determination of ***W***, we introduced a modified cross-validation function and choose the ***W*** in 2.1.3.

#### 2.1.2 Estimation of low-rank correlation matrices based on MM algorithm

The MM algorithm for estimating a low-rank correlation matrix proposed by Simon and Abell [2010] is explained. To estimate ***Y*** in the closed form under the constraint (2), the quadratic optimization problem for ***Y*** must be converted to a linear optimization one. Using the linear function, we can derive the updated formula combined with the Lagrange multiplier in the closed form. Let 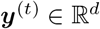 be the parameter of *t* step in the algorithm of the optimization problem, and *g*(***y***|***y***^(*t*)^) be a real function 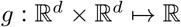. If *g*(***y***|***y***^(*t*)^) satisfy the following conditions such that

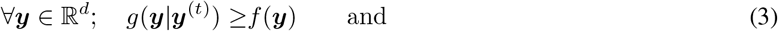

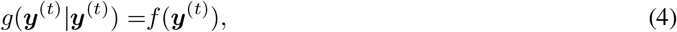

*g*(***y***|***y***^(*t*)^) is defined as the majorizing function of *f*(***y***) at the point ***y***^(*t*)^, where 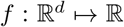 is the original function. Simply put, to estimate the parameters in the MM algorithm, *g*(***y***|***y***^(*t*)^), not *f*(***y***), should be minimized. In several situations, *g*(***y***|***y***^(*t*)^) is expected to easily minimize the value. For more details regarding the MM algorithm, see Hunter and Lange [2004].

Before deriving the majorizing function, the objective function (1) can be redescribed as follows:

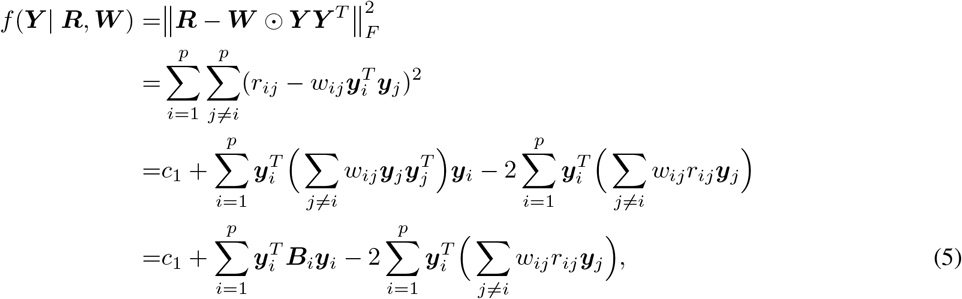

where 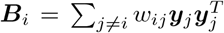 and *c*_1_ are constants. Here, the parameter estimation of ***Y*** is conducted by ***y***_*i*_. The corresponding part of Eq. (5) and the majorizing function can be described as follows:

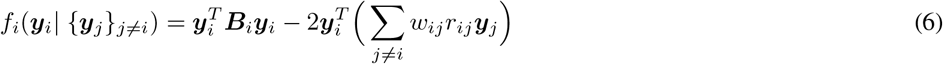

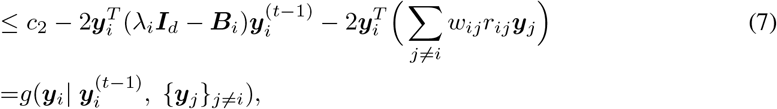

where 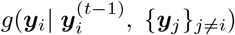 represents the majorizing function of Eq.(5), *c*_2_ is constant. ***I***_*d*_ is *d* × *d* identity matrix, *λ*_*i*_ is the maximum eigenvalue of ***B***_*i*_, and 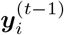 is ***y***_*i*_ of (*t* − 1) step in the algorithm. Here, the inequality of Eq. (7) is satisfied because ***B***_*i*_ − *λ*_*i*_***I***_*d*_ is negative semi-definite. In fact, if 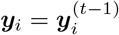, Eq.(6) and Eq.(7) becomes equal.

Using the Lagrange multiplier method and Eq. (7), the updated formula of ***y***_*i*_ is derived as follows:

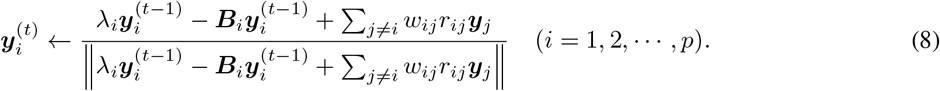

##### Algorithm 1 Algorithm for estimating the low-rank correlation matrix

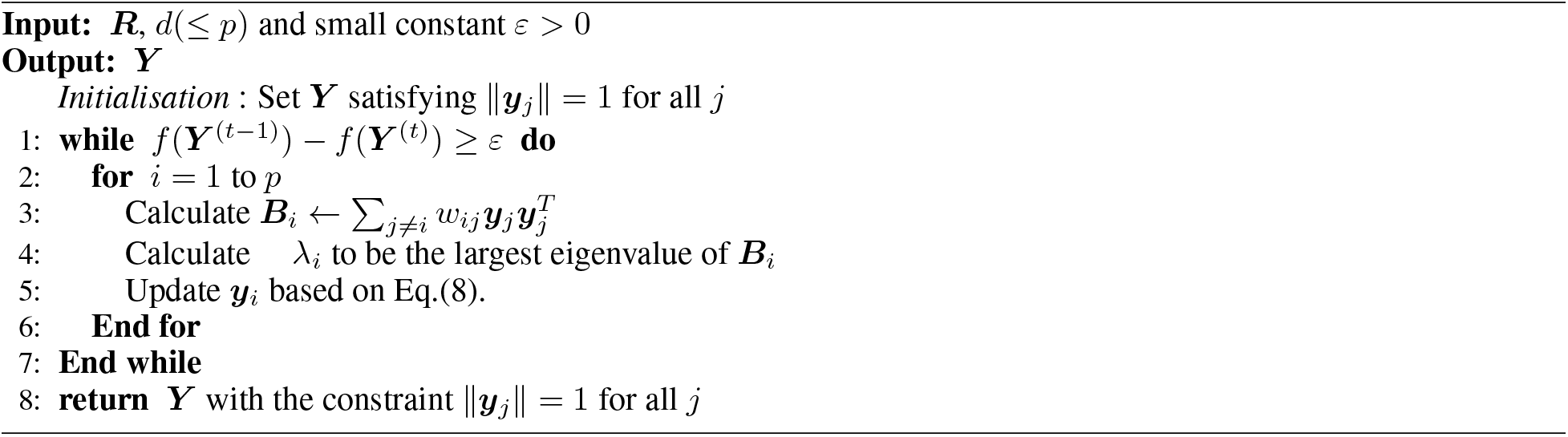

#### 2.1.3 Proposed Algorithm of Cross Validation to determine hard thresholds

In the proposed approach, we adopt hard thresholding to estimate the sparse low-rank correlation matrix. To determine the threshold values, we introduce a cross-validation function based on Bickel and Levina [2008a]. The purpose of this approach is quite simple, that is, to determine the threshold values related to sparse estimation by considering the corresponding rank.

Let *h*(*α*) ∈ (−1, 1) be a threshold value of sample correlation coefficients corresponding to the *α* percentile of correlations, where *α* ∈ [0, 1] is the percentage point. From that, by the setting percentile point *α*, the corresponding threshold value *h*(*α*) is fixed. For a correlation *r*_*ij*_ ∈ [−1, 1], the function 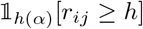 is defined as 1 if *r*_*ij*_ ≥ *h*(*α*), otherwise 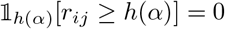. Using them, proportional threshold operator is defined as

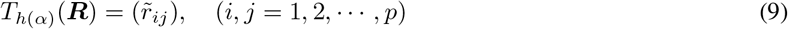

where

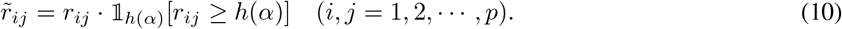

For example, the proportional threshold operator is used in the domain of neural science [van den Heuvel et al., 2017]. Let 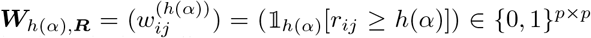 for correlation matrix ***R***. By using ***W***_*h*(*α*)*,**R***_, Eq.(9) can be described as follows:

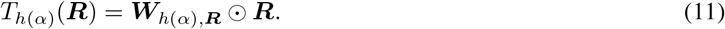

Here, Eq.(10) is modified for the original function of Bickel and Levina [2008a]. Originally, 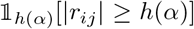 is used, however, we only focus on higher correlation coefficients and not on negative correlation coefficients. Using modifications, it becomes easy to interpret the results.

To estimate a sparse a low-rank correlation matrix, we introduce the modified proportional threshold operator based on Eq.(9) because the interpretation of the proportional threshold is quite simple. Given a *α* percentile, the rank *d* and correlation matrix ***R***, the modified threshold operator is defined as follows:

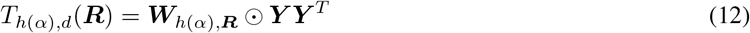

where ***Y*** is the estimated correlation matrix with rank *d* such as minimizing *f* (***Y*|*R***, ***W***_*h*(*α*)*,**R***_). Eq. (12) is different from Eq. (12) in using a low rank correlation matrix, although ***W***_*h*(*α*)*,**R***_ is calculated from the original correlation matrix, not from a low rank correlation matrix. For the choice of the threshold value *h*(*α*), a cross-validation was introduced (e.g., Bickel and Levina [2008a], Jiang [2013]). The cross-validation procedure for the estimation of *h*(*α*) consists of three steps, as shown in Figure1. First, original multivariate data 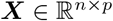 is split into two groups randomly such as 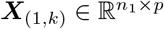 and 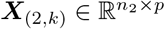, where *n*_1_ = *n* −⌊*n/* log *n*⌋, *n*_2_ = ⌊*n/* log *n*⌋ and *k* represents the index of the number of iterations for cross validation, ⌊·⌋ represents floor function. For *n*_1_ and *n*_2_, Bickel and Levina [2008a] determines both *n*_1_ and *n*_2_ from the perspective of theory. Second, correlation matrices for both ***X***_(1*,k*)_ and ***X***_(2*,k*)_ are calculated as ***R***_(1*,k*)_ and ***R***_(2*,k*)_, respectively. Third, the correlation matrix with rank *d*, ***Y***, is estimated such as minimizing 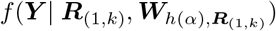 with constraint (2). Forth, for fixed *h*(*α*), the procedure from the first step to the third step is repeated *K* times and the proposed cross-validation function is calculated as follows.

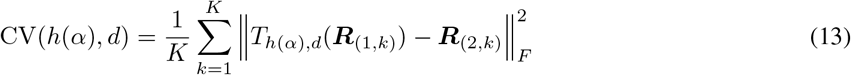

where *K* is the number of iterations for the cross-validation. Among the candidates of threshold values, *h*(*α*) is selected as the value such that the expression in Eq.(13) is minimized. The algorithm for the cross-validation is presented in **Algorithm 2**.

##### Algorithm 2 Algorithm of Cross-validation for tuning proportional thresholds

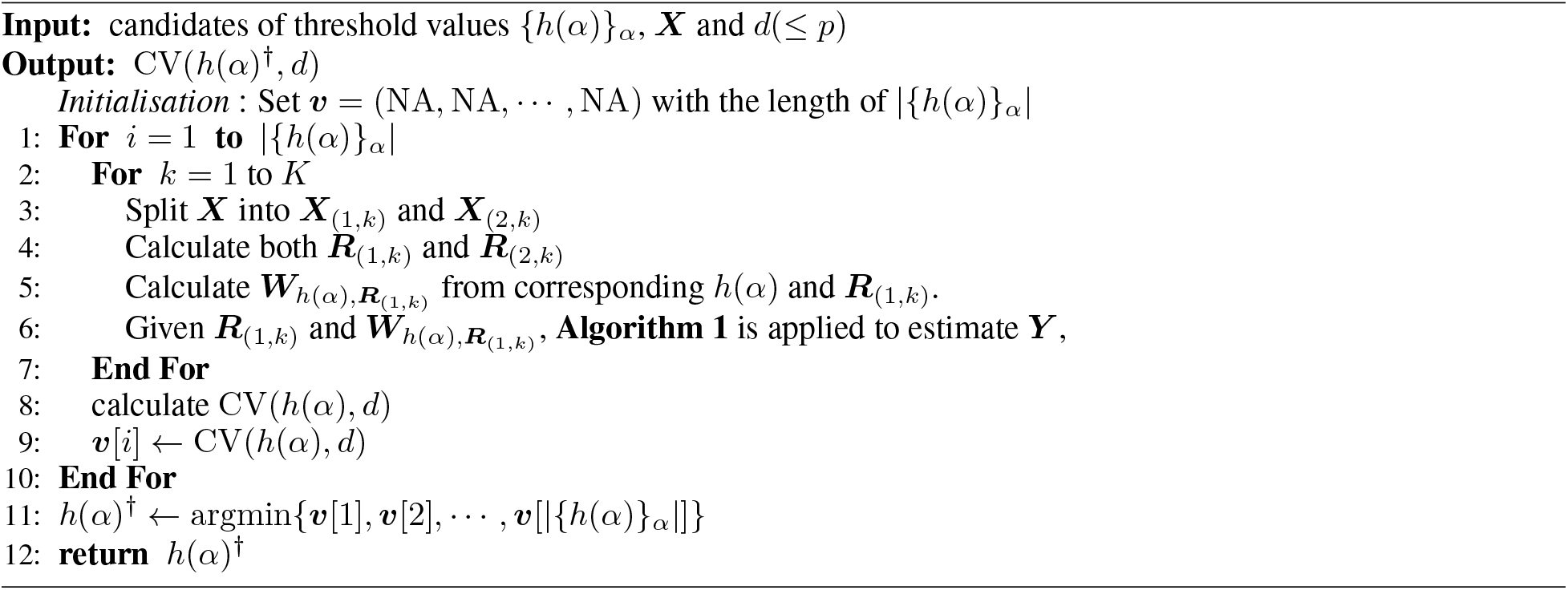

**Figure 1:**
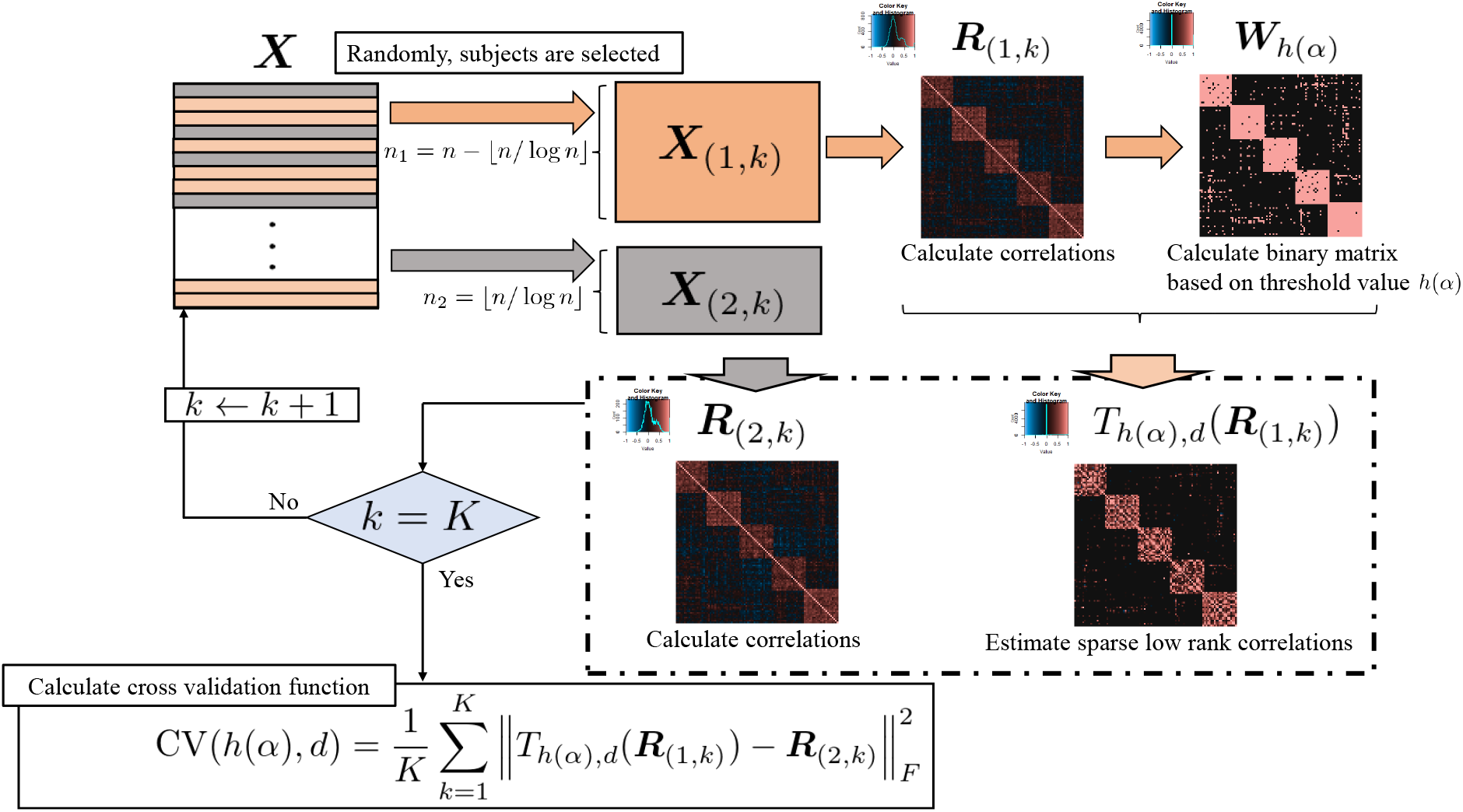
Framework of the proposed cross validation

Finally, *h*(*α*)^†^, corresponding to the minimum value of Eq.(13) among the candidate threshold values is selected and 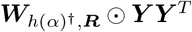 is estimated based on Eq.(1).

### 2.2 Numerical Simulation and Real example

This section presents a numerical simulation to evaluate the proposed approach. The numerical simulation was conducted based on Cui et al. [2016] with some modification. Practically, the size of numerical data is matched to that of the real data example in 2.2.2. In addition, we present a real example of applying the proposed method to a micro-array gene expression dataset from Khan et al. [2001].

#### 2.2.1 Simulation design of numerical simulation

In this subsection, the simulation design is presented. The framework of the numerical simulation consists of three steps. First, artificial data with a true correlation matrix are generated. Second, sparse low-rank correlation matrices are estimated using two methods, including the proposed method. In addition, a sample correlation matrix and sparse correlation matrix based on threshold also apply. Third, using several evaluation indices, these estimated correlation matrices are evaluated and their performances are compared.

In this simulation, three kinds of correlation models are used. Let *I* and *J* be a set of indices for the rows and columns of correlation matrices, respectively. In addition, *I*_*k*_ and *J*_*k*_ are defined as follows:

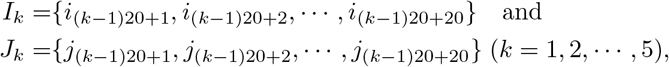

where *i*_*o*_ and *j*_*o*_ (*o* = 1, 2, · · ·, 100) indicates the number of rows and columns, respectively. Using these notations, three true correlation models 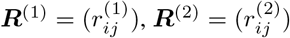, and 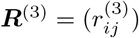 are set as

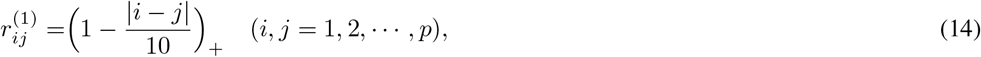

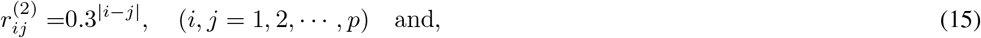

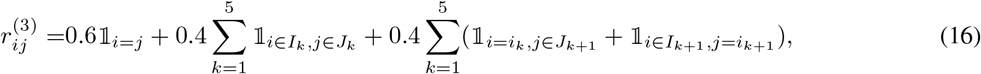

respectively, where *i*_*k*_ and *j*_*k*_ are the maximum number of indices of *I*_*k*_ and *J*_*k*_, respectively, and 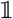 represents indicator function. The models for Eq.(14) and Eq.(15) are called sparse models, while the model for Eq.(16) is called a non-sparse model by Cui et al. [2016]. The models for Eq.(14) and Eq.(15) are used in Bickel and Levina [2008a], Xue et al. [2012], and Rothman [2012]; for these, see Figure 2. These artificial data are generated as 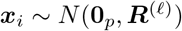 (*i* = 1, 2, · · ·, *n*; *ℓ* = 1, 2, 3), where **0**_*p*_ is a zero vector with a length of *p*. In this simulation, we set *p* = 100 and the number of cross-validations *K* = 5. In this simulation, there are several kinds of scenarios. For the number of scenarios in the estimation of sparse low-rank correlation matrices, there are 2(setting1) × 3(setting2) × 3(setting3) × 2(setting4: proposal and tandem) = 36 patterns. In addition to that, for the number of scenarios in the estimation of sparse correlation matrices without low-rank approximation, there are 2(setting1) × *three*(setting3) × 2(setting4: proposal and tandem) = 12 patterns. Simply, there are 48 patterns in this numerical simulation. In each pattern, artificial data is generated 100 times and evaluated using several indices. In addition, Both the proposed approach and tandem approach run from a random start 50 times, and the best solution is selected. For ***R***_1_ and ***R***_2_, the candidates of *α* are set as 0.66 to 0.86 in steps of 0.02. However, for ***R***_1_, the candidates of *α* are set from 0.66 to 0.82 in steps of 0.02.

**Figure 2:**
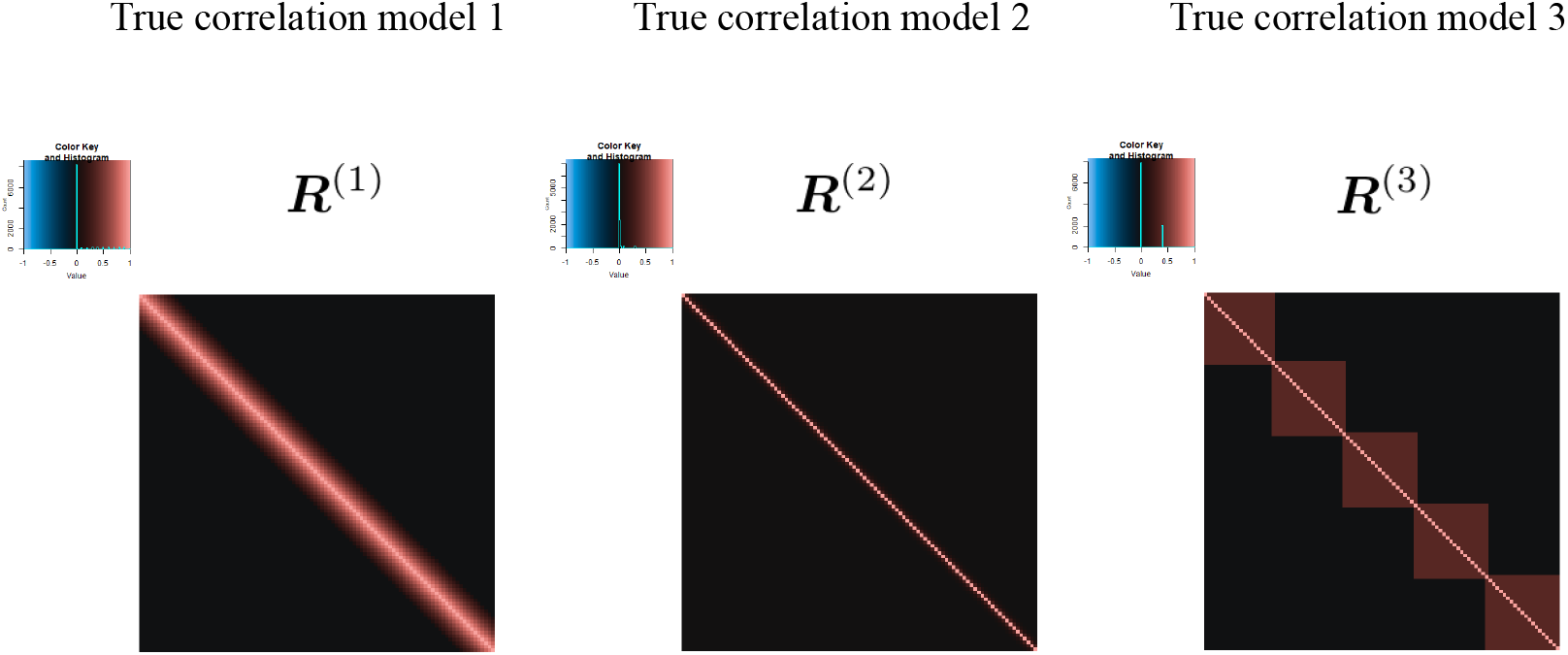
True correlation models

Next, the settings of the numerical simulation are presented. For the summary, see Table 1. Setting one was set to evaluate the effects of the number of subjects. If the number of subjects is smaller, the estimated sparse low-rank correlation is expected to be unstable. To evaluate the effect of rank, setting two was set. When a smaller rank is set, the variance between the estimated sparse low-rank correlation coefficients becomes larger. Therefore, it becomes easy to interpret the results. Next, as explained in Eq.(14), Eq.(15) and Eq.(16), there are three levels in setting three. Finally, in setting four, we set four methods: the proposed approach, tandem approach, sample correlation matrix calculation, and sparse correlation matrix estimation based on threshold value [Jiang, 2013] with modifications. The purpose of both the proposed approach and the tandem approach is to estimate a sparse low-rank correlation matrix. In the tandem approach, estimation of the low-rank correlation matrix is the same as that of the proposed approach; however, the determination way of proportional threshold is different from Eq.(13). In contrast to Eq.(13), the proportional threshold is selected using the method of Bickel and Levina [2008a] and Jiang [2013] with modification, which does not consider the features of the low-rank correlation matrix. Here, in the tandem approach, we use Eq.(10) as a threshold function. Therefore, in the tandem approach, given the corresponding ***W***_*h*(*α*)_ and correlation matrix ***R***, a sparse low-rank correlation matrix is estimated based on the optimization problem of Eq.(1). To estimate the sparse correlation matrix without dimensional reduction, Eq.(10) is used as the threshold function: although 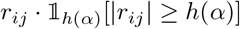 was used in Jiang [2013]. In this paper, we follow the approach without dimensional reduction, Jiang (2013) with modifications.

**Table 1:** Settings of numerical simulation

| Setting name | Levels | Description |
| --- | --- | --- |
| Setting 1: The number of subjects | 2 | $n = 50, 75$ |
| Setting 2: Rank | 3 | $d = 2, 3, 5$ |
| Setting 3: True correlation model | 3 | Eq.(14), Eq.(15) and Eq.(16) |
| Setting 4: Methods | 4 | proposed approach, tandem approach<br>sample correlation, and Jiang (2013) with modification |

Likewise as the approach pursued in Cui et al. [2016], we adopt four evaluation indices. To evaluate the fitting between estimated sparse low-rank correlation matrix and true correlation matrix, the average relative error of the Frobenius norm (F-norm) and of a spectral norm (S-norm) are adopted as follows:

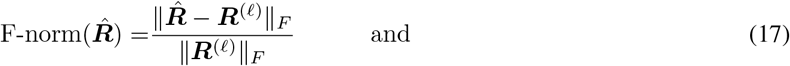

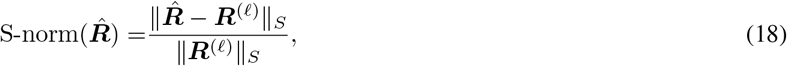

respectively, where ∥ · ∥_*S*_ indicates spectral norm, 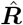 is an estimator of the sparse low-rank correlation matrix, and 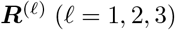 is the true correlation matrix corresponding to Eq.(14), Eq.(15) and Eq.(16), respectively. In addition, to evaluate the results on sparseness, the true positive rate (TPR) and false-positive rate (FPR) are defined as follows:

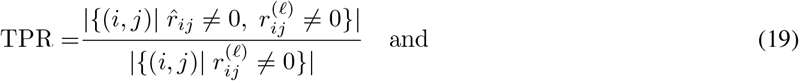

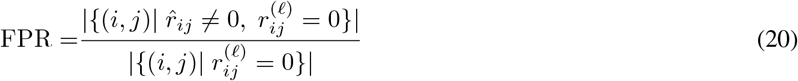

where | · | indicates the cardinality of a set, 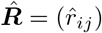, and 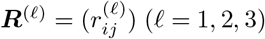.

#### 2.2.2 Application of microarray gene expression dataset

Here, we present the results of applying both the proposed approach and the tandem approach to the microarray gene expression dataset in Khan et al. [2001]. The purpose of this real application is to evaluate the differences between two classes of genes on the results of estimating sparse low-rank correlation matrices. Concretely, in this real example, true correlation coefficients between classes are assumed to be zero, and the FPR of the proposed approach is compared with that of the tandem approach.

In Rothman et al. [2009], the same dataset was used as an application of their method. Specifically, the dataset provided by the R package “MADE4” [Culhane et al., 2005] is used in this example. The dataset includes 64 training sample and 306 genes. In addition, there are four types of small round blue cell tumors of childhood (SRBCT), such as neuroblastoma (NB), rhabdomyosarcoma(RMS), Burkitt lymphoma, a subset of non-Hodgkin lymphoma (BL) and the Ewing family of tumors (EWS). Simply, there are four sample classes in this dataset. As was done in Rothman et al. [2009], these genes are classified into two classes: “informative” and “noninformative,” where genes belonging to “informative” have information to discriminate four classes and those belonging to “noninformative” do not.

Next, to construct “informative” class and “noninformative” class, F statistics is calculated for each gene as follows:

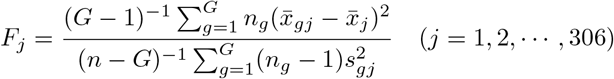

where *G* indicates the number of classes, such as NB, RMS, BL, and EWS, *n*_*g*_ is the number of subjects belonging to class *g*, and 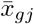 is mean of class *g* for gene *j*, and 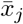 is the mean of gene *j*, *s*_*gj*_ is the sample variance of the class *g* for gene *j*. Here, if *F*_*j*_ is relatively higher, gene *j* is considered as “informative” because the corresponding *j* tends to include information such that each class is discriminated. From the calculated *F*_*j*_, the top 40 genes and bottom 60 genes are set as “informative” class and “noninformative” class, respectively. Then, the correlation matrix for 100 genes was calculated and set as input data. See Figure 3.

**Figure 3:**
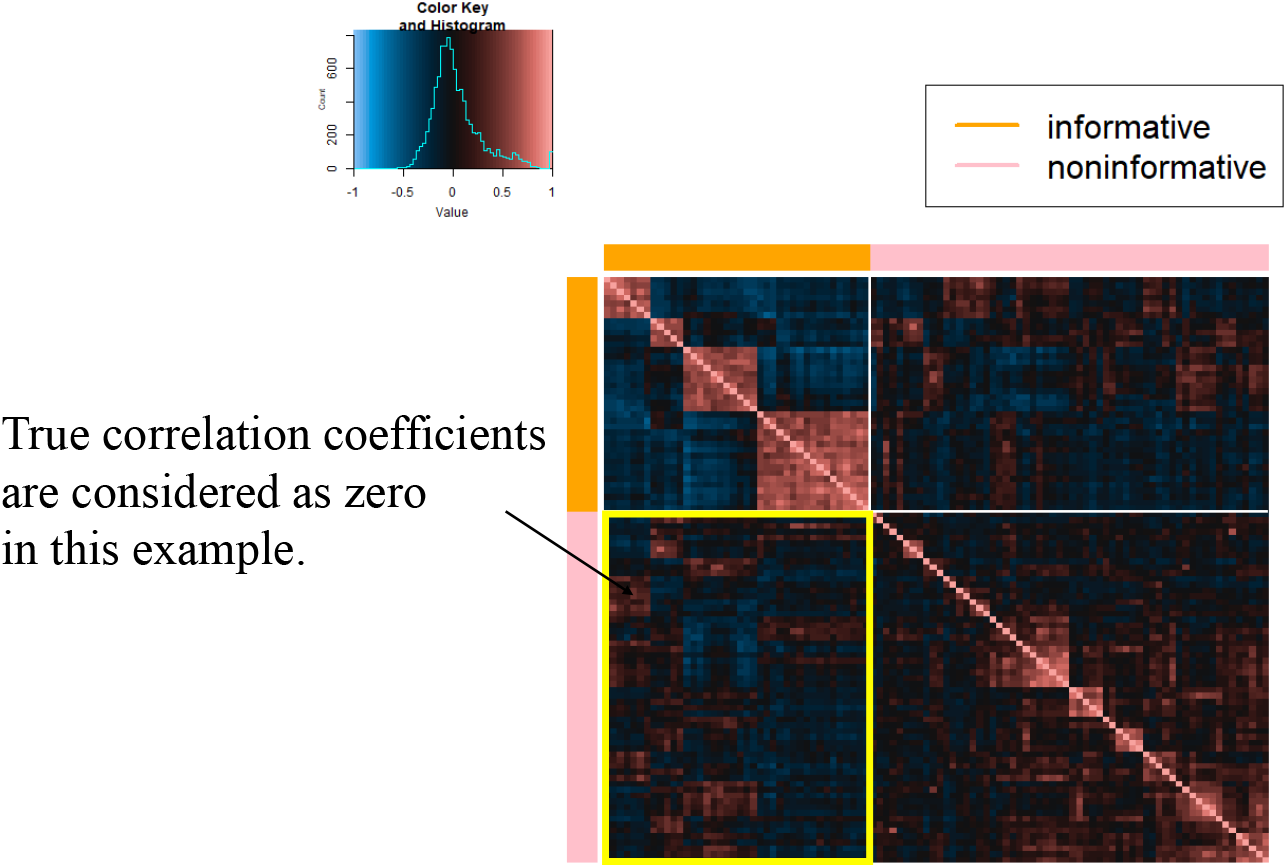
Sample correlation matrix among selected 100 genes

To compare the results of the proposed approach with those of the tandem approach, the FPR is calculated. For the tandem approach, see “2.2.1 Simulation design of numerical simulation". In this application, true correlations between genes belonging to “informative” class and gene belonging to “noninformative” class is considered as 0. Therefore, the denominator of the FPR is set to 2 × 40 × 60 = 4800. For TPR, it is difficult to determine the true structure because correlations within each class are not necessarily non-zero. For the rank, we set 2, 3, and 5. The candidates of *α* for determining the threshold value are set as 0.50 to 0.83 in steps of 0.01 for both approaches, and these algorithms start from 50 different initial parameters. In addition, as was done for the numerical simulation, both the sample correlation matrix and Jiang (2013) with modifications are also employed.

## 3 Results

This section presents the results of the numerical simulation and real application.

### 3.1 Simulation result

In this subsection, we present the simulation results by the true correlation models. Table 2, Table 3 and Table 4 indicate the FPRs and TPRs for applying ***R***^(1)^, ***R***^(2)^, and ***R***^(3)^, respectively. Each cell indicates mean of these indices. Here, ***R***^(2)^ is a non-sparse correlation matrix and therefore, FPR cannot be calculated, and both the TPR and FPR of the sample correlation matrix cannot be calculated because the sample correlation is not a sparse matrix. From the results of the numerical simulation, the FPRs of the proposed approach was the lowest among those of all the methods in all situations. while the TPRs of the proposed approach tended to be inferior to those of the other approaches. Simply, the proposed approach makes it a sparser low-rank correlation matrix compared to the tandem approach when a smaller rank is used.

**Table 2:**
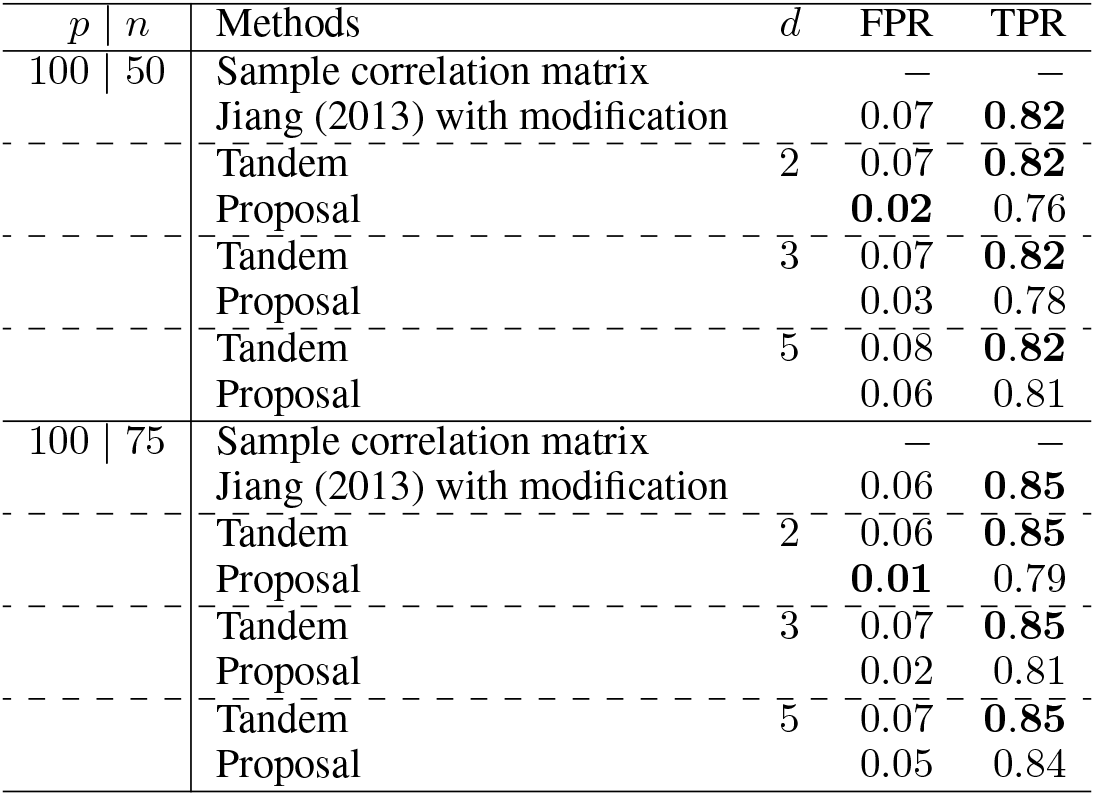
Results of FPRs and TPRs for ***R***^(1)^ and each value indicate the mean.

**Table 3:**
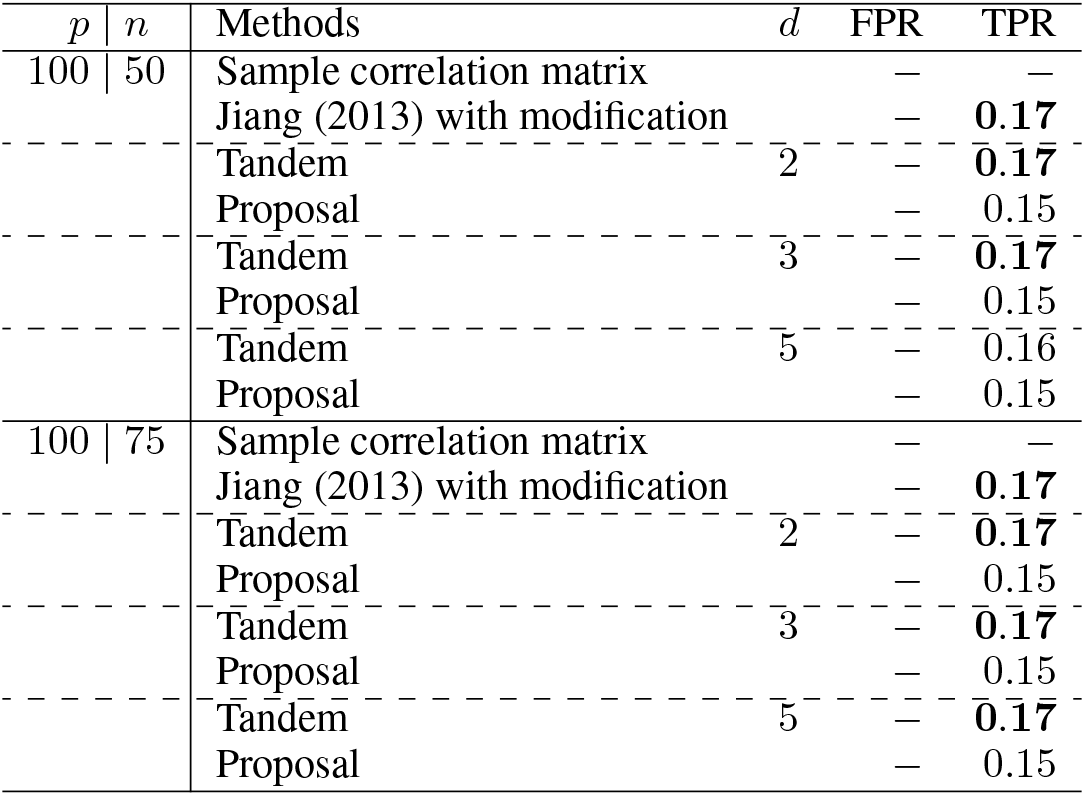
Results of FPRs and TPRs for ***R***^(2)^ and each value indicate the mean.

**Table 4:**
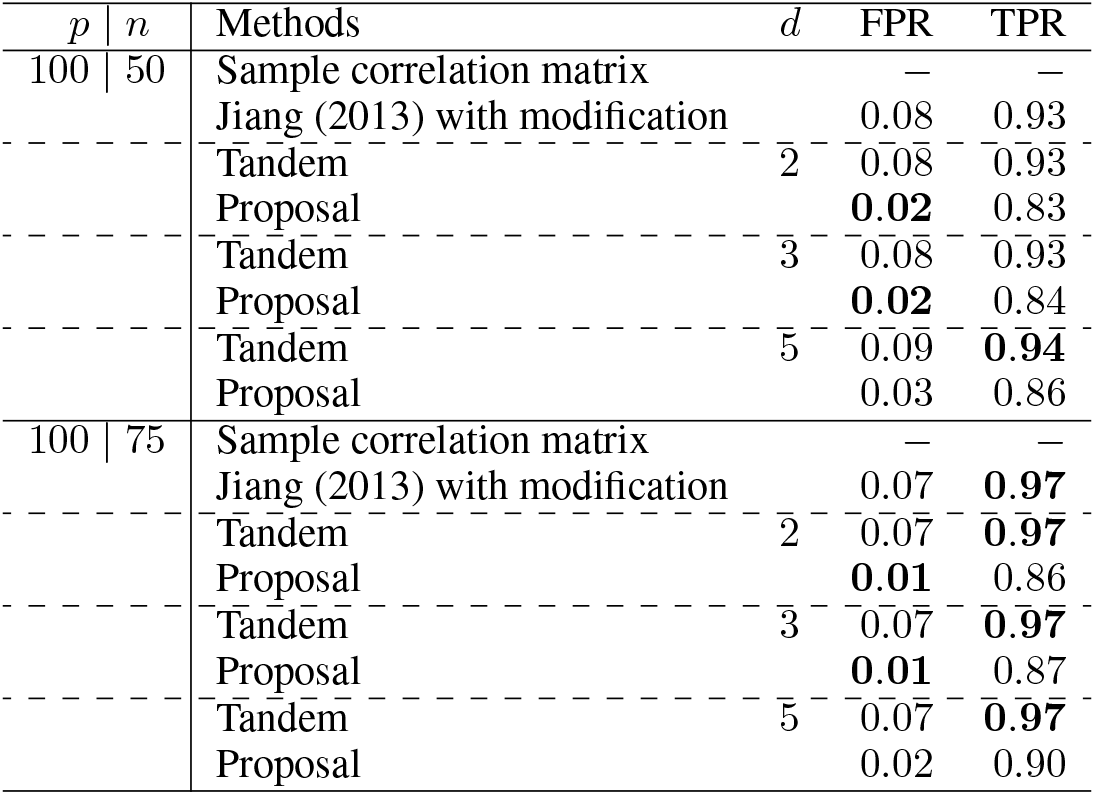
Results of FPRs and TPRs for ***R***^(3)^ and each value indicate the mean.

For the relative error of F-norm, Figure 4, Figure 5 and Figure 6 indicates the results of applying these methods to ***R***^(1)^, ***R***^(2)^, and ***R***^(3)^, respectively. Hence, the median of the proposed approach was lower than that of the tandem approach for each pattern. In addition, the interquartile range of the proposed approach was smaller than that of the tandem approach in each pattern. Therefore, we confirmed that the results of the proposed approach are effective and stable compared to those of the tandem approach. As rank is set as larger, The results of both the proposed approaches become lower and close to those of Jiang (2013) with modifications in all situations. Among these methods, the result of Jiang (2013) with modifications is the best for the relative error of F-norm. However, it is natural things from the properties of low-rank approximation. As was done for F-norm, those of S-norm for ***R***^(1)^, ***R***^(2)^ and ***R***^(3)^ are shown in Figures 7, 8 and 9, respectively. The tendency of the results for S-norm is quite similar to that for F-norm. From the results for F-norm, we observe that the result of the proposed approach with rank 5 is quite close to that of Jiang (2013) with modification.

**Figure 4:**
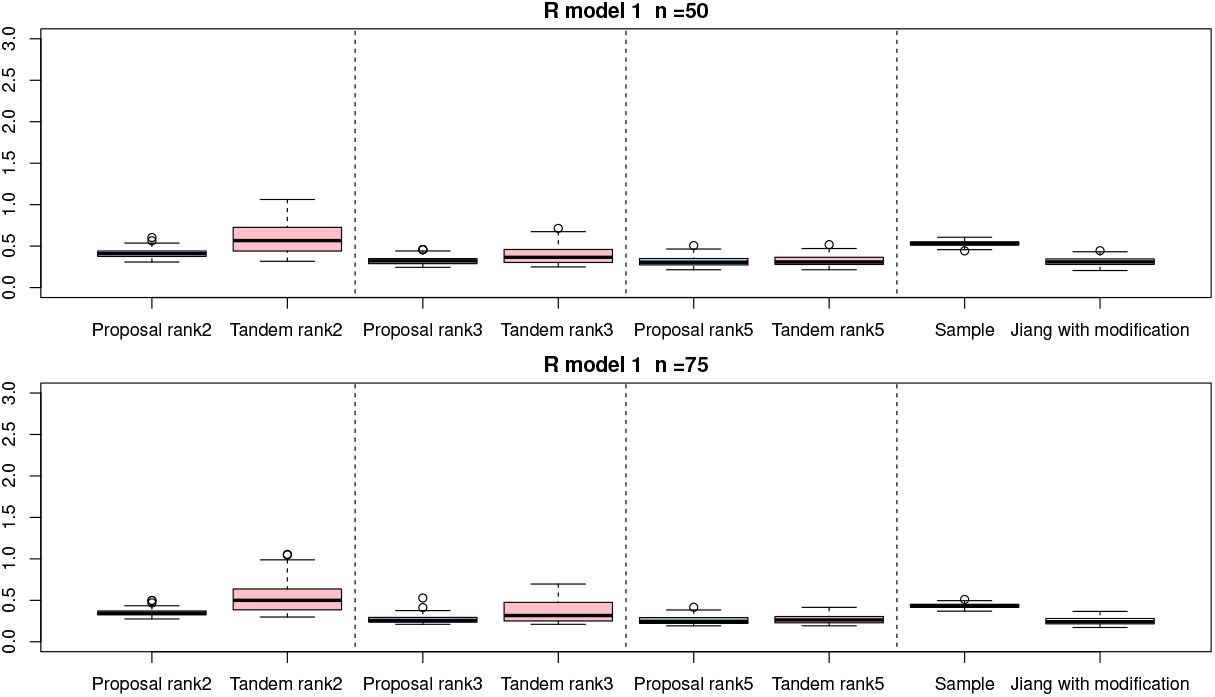
Relative errors of F-norm for ***R***^(1)^ with *n* = 50 and *n* = 75; the vertical axis indicate the results of relative errors of F-norm.

**Figure 5:**
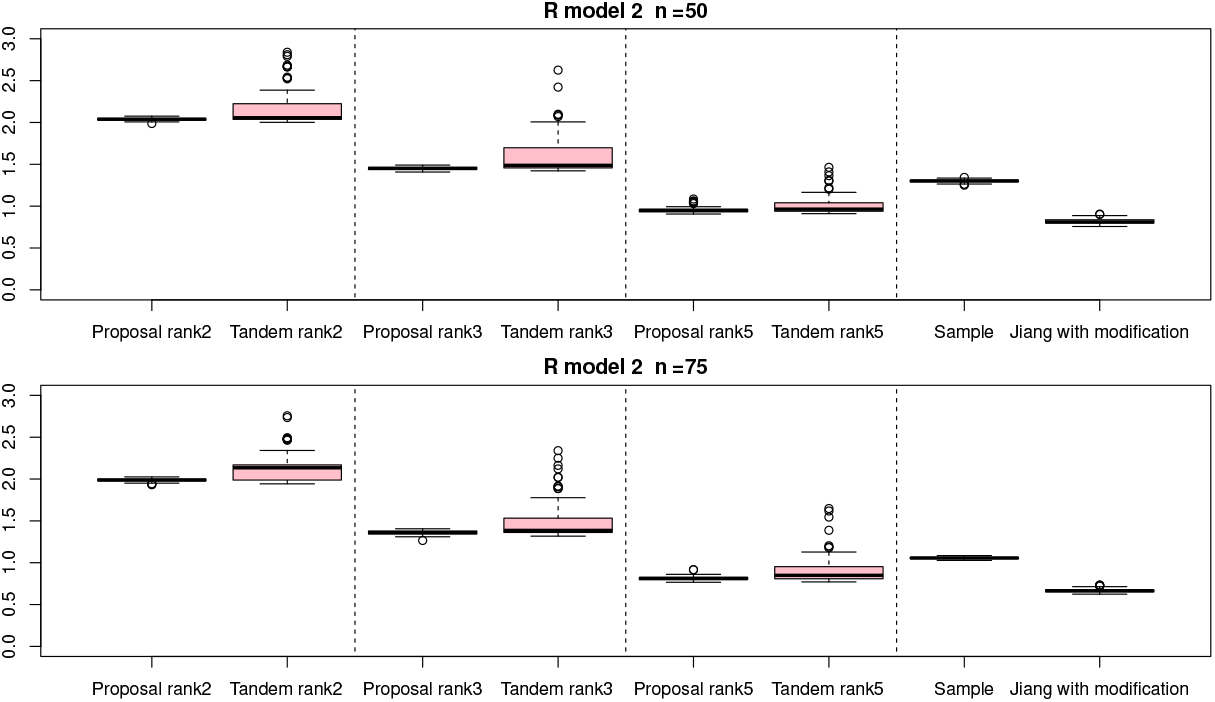
Relative errors of F-norm for ***R***^(2)^ with *n* = 50 and *n* = 75; the vertical axis indicate the results of relative errors of F-norm.

**Figure 6:**
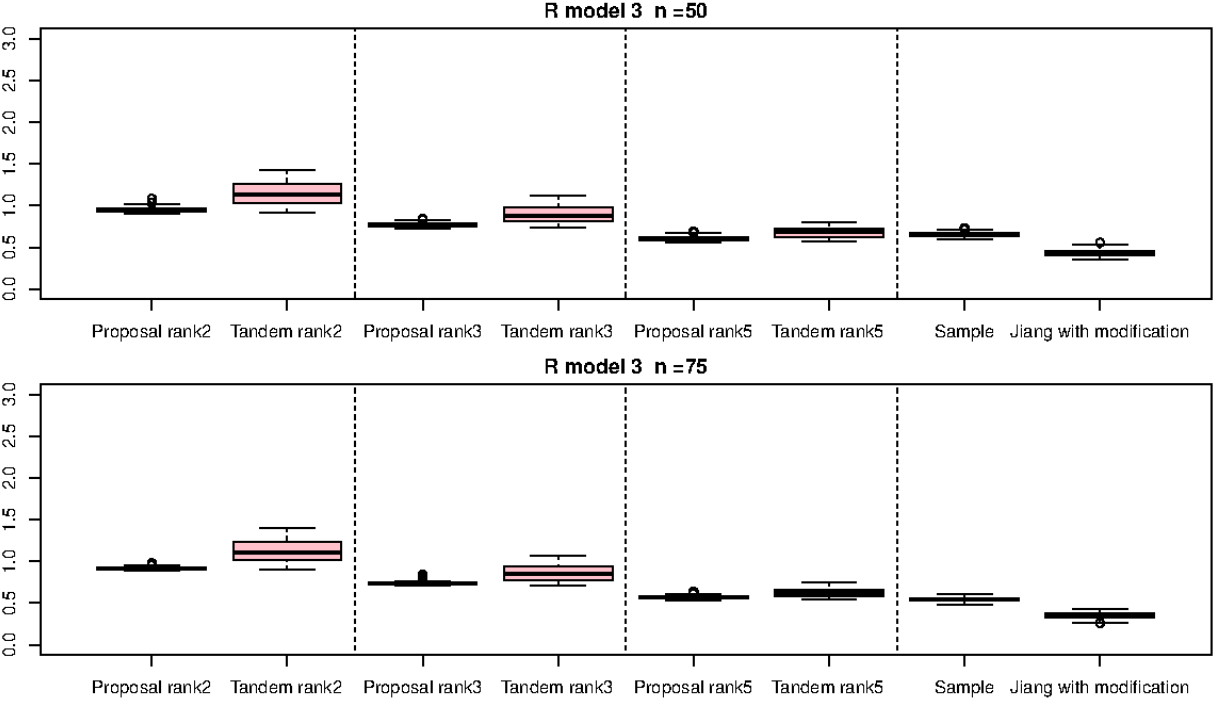
Relative errors of F-norm for ***R***^(3)^ with *n* = 50 and *n* = 75; the vertical axis indicate the results of relative errors of F-norm.

**Figure 7:**
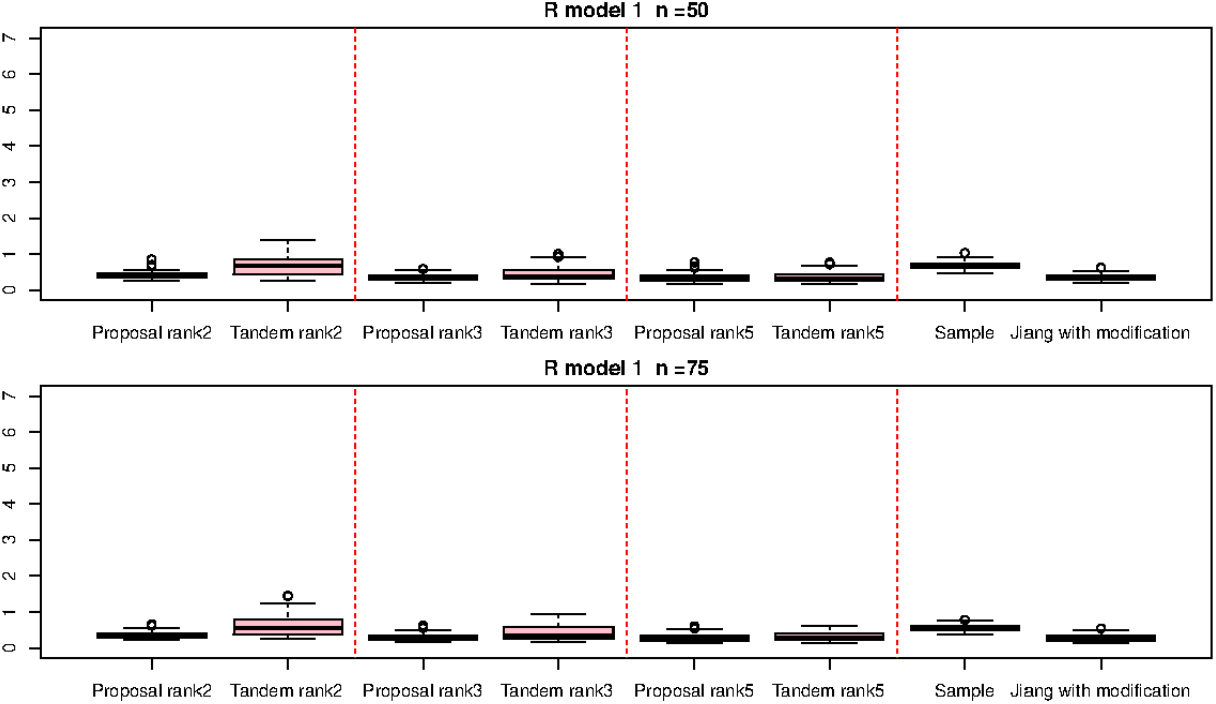
Relative errors of S-norm for ***R***^(1)^ with *n* = 50 and *n* = 75; the vertical axis indicate the results of relative errors of S-norm.

**Figure 8:**
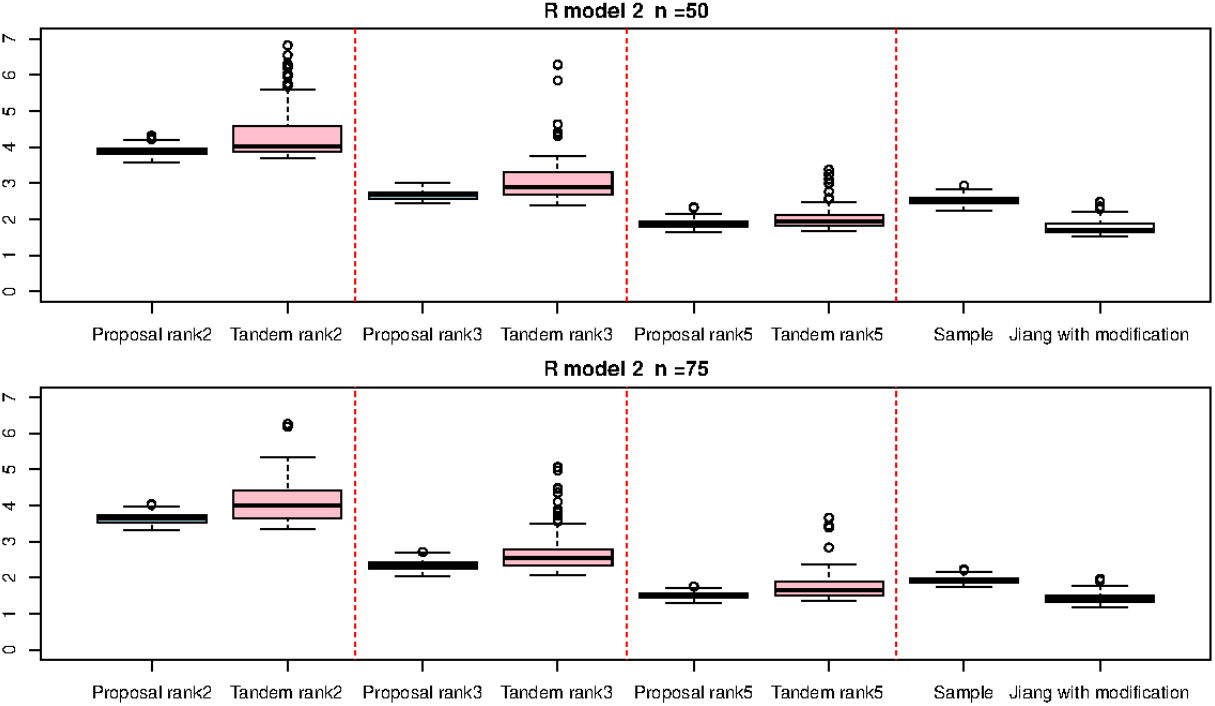
Relative errors of S-norm for ***R***^(2)^ with *n* = 50 and *n* = 75; the vertical axis indicate the results of relative errors of S-norm.

**Figure 9:**
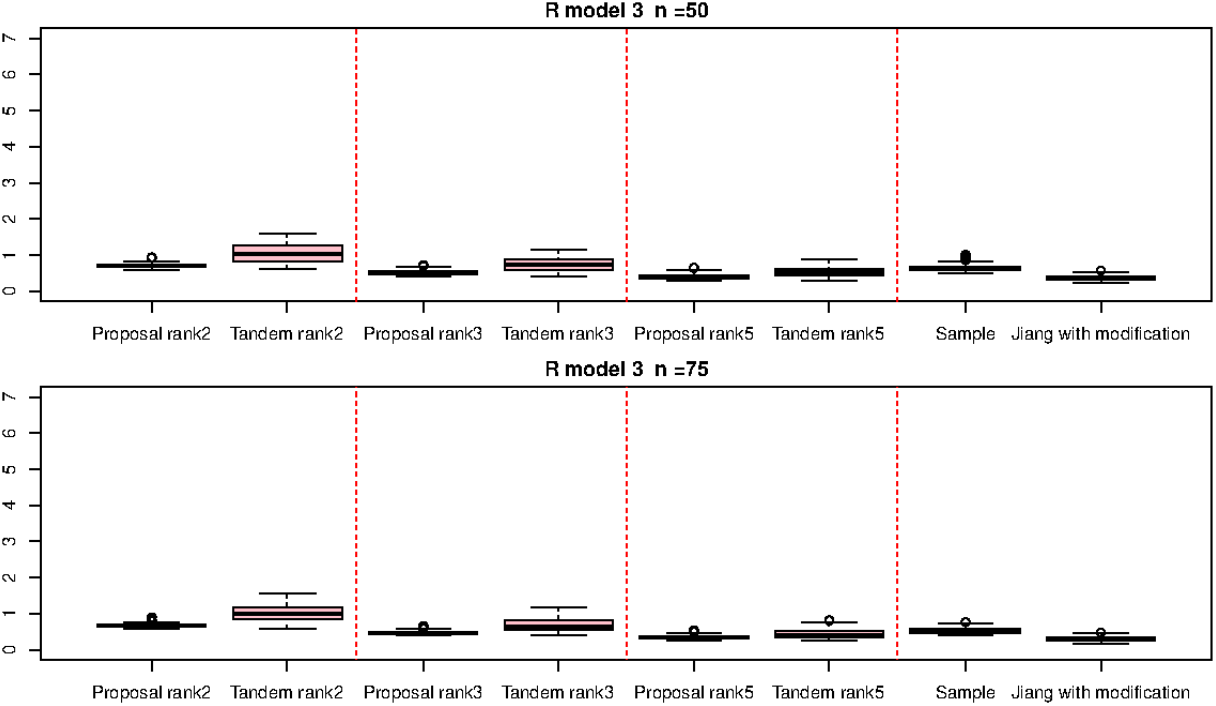
Relative errors of S-norm for ***R***^(3)^ with *n* = 50 and *n* = 75; the vertical axis indicate the results of relative errors of S-norm.

For the estimated correlation matrices, Figure 10, Figure 11 and Figure 12 correspond to true correlation model one, true correlation model 2, and true correlation model 3 with *n* = 50, respectively. As the same way, Figure 13, Figure 14 and Figure 15 correspond to true correlation model one, true correlation model 2, and true correlation model 3 with *n* = 75, respectively. From Figures 10, 13, 12 and 15, we found that the estimated correlation matrices of the proposed approach tend to estimate zero correctly compared to those of the tandem approach. Especially, the tendency can be confirmed when the rank is set as lower, visually. In addition, rank is set larger, estimated correlation matrices tend to be close to the results of Jiang (2013) with modifications.

**Figure 10:**
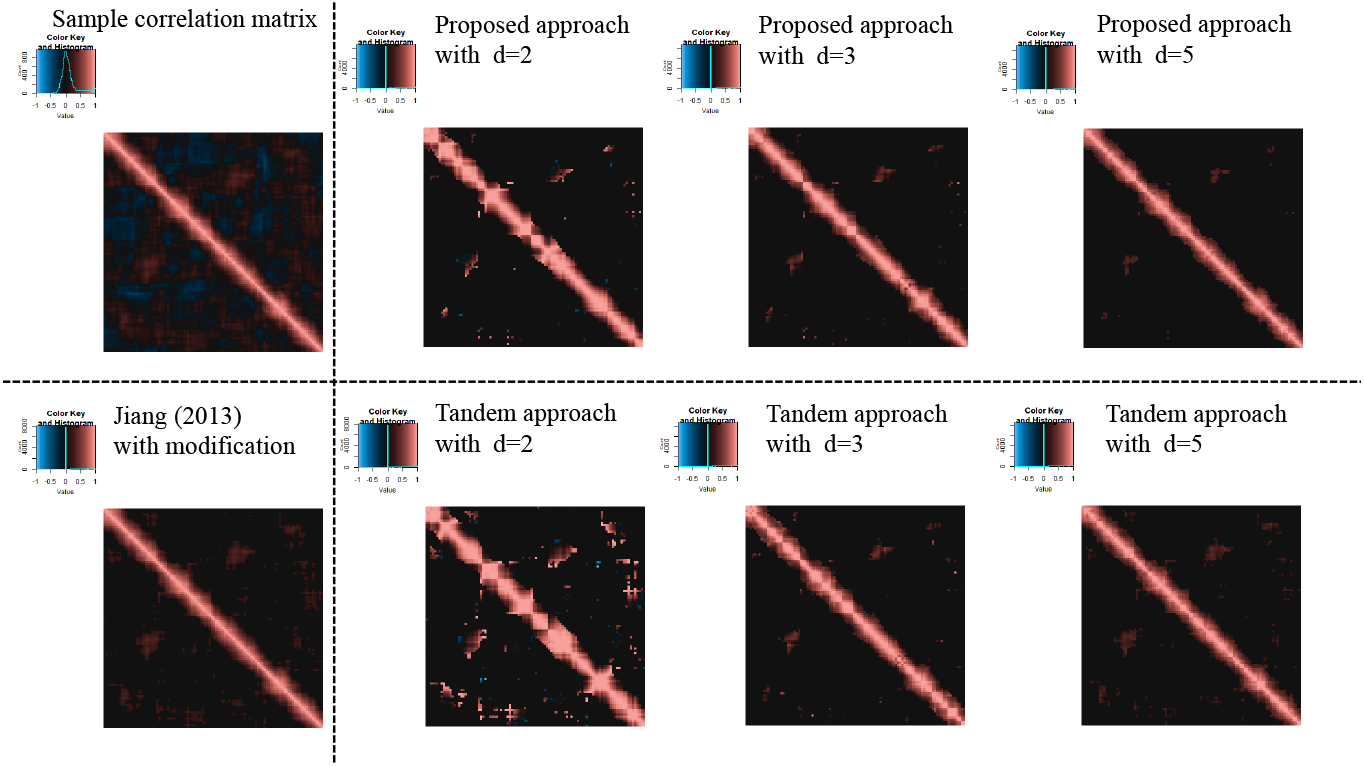
Examples of estimated correlation matrices for true correlation model 1 (*n* = 50)

**Figure 11:**
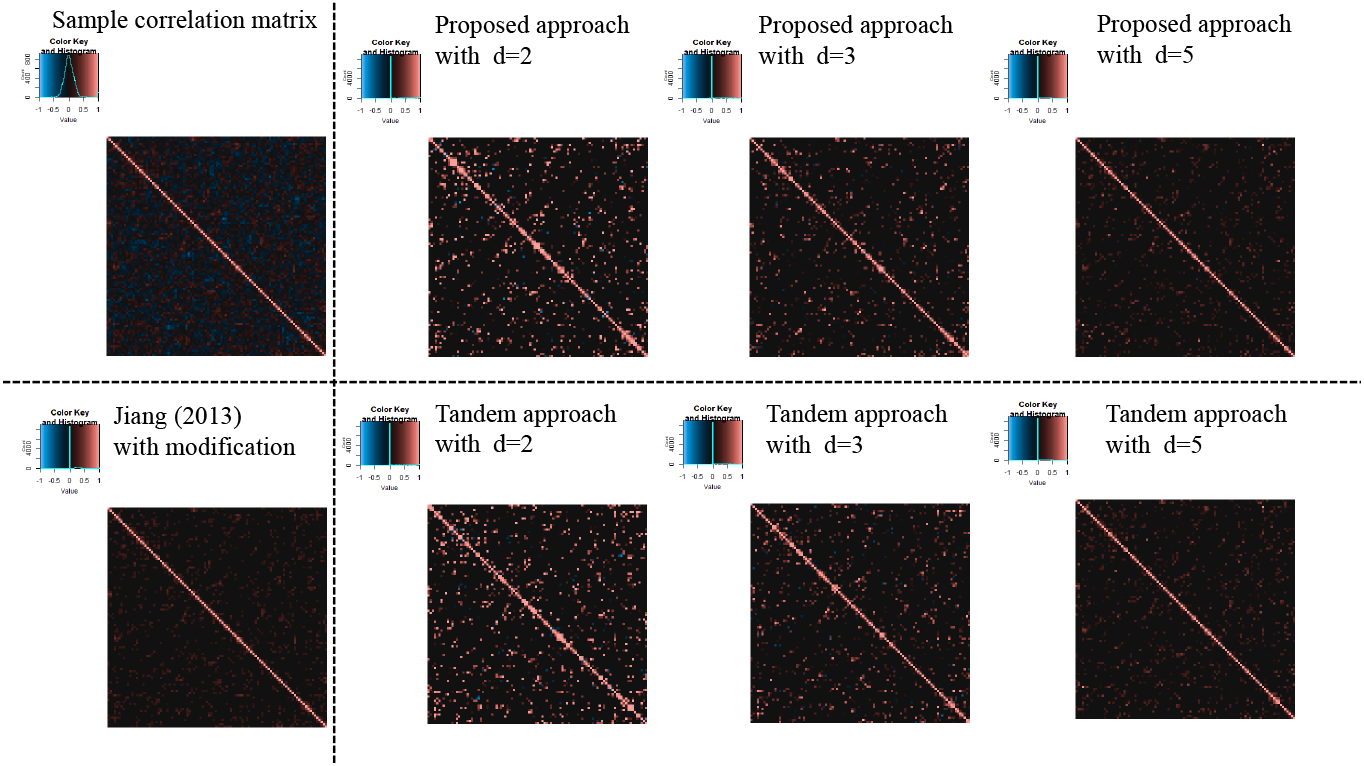
Examples of estimated correlation matrices for the true correlation model 2 (*n* = 50)

**Figure 12:**
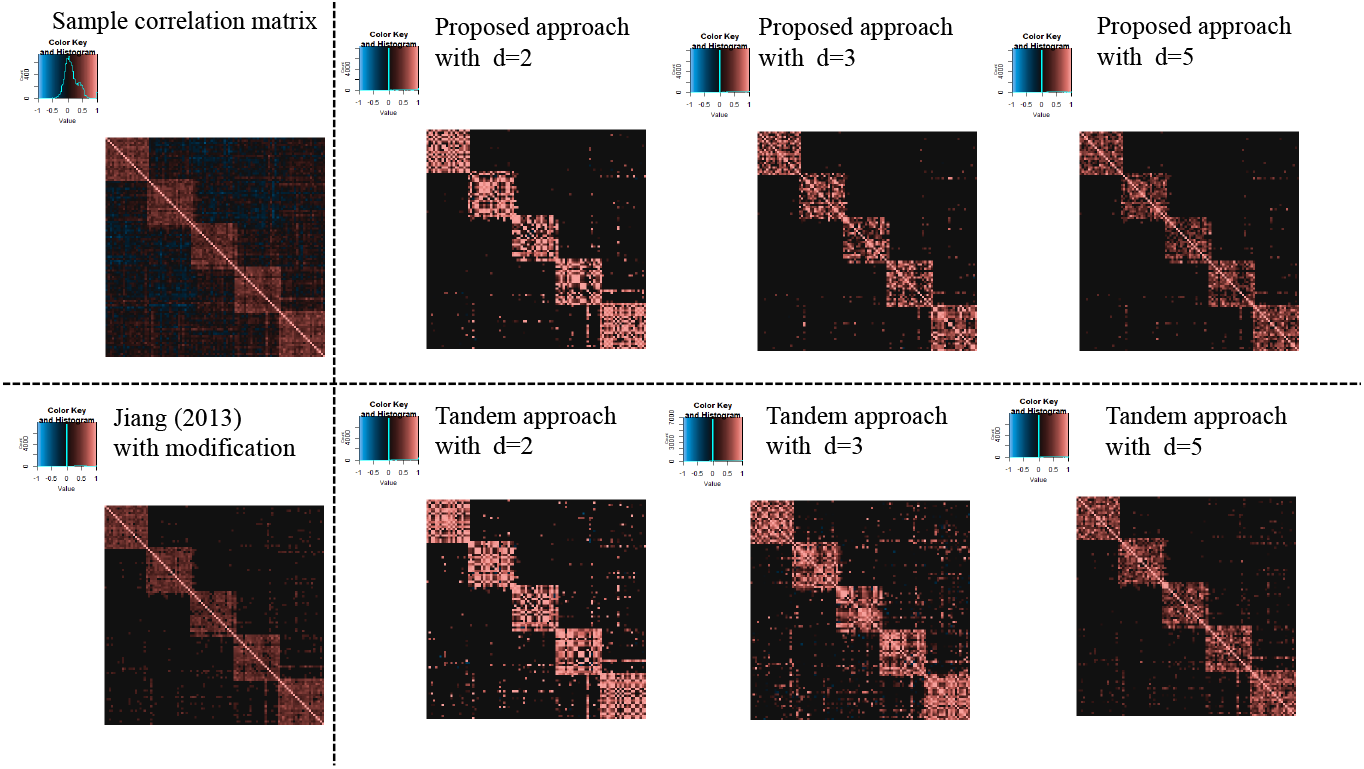
Examples of estimated correlation matrices for the true correlation model 3 (*n* = 50)

**Figure 13:**
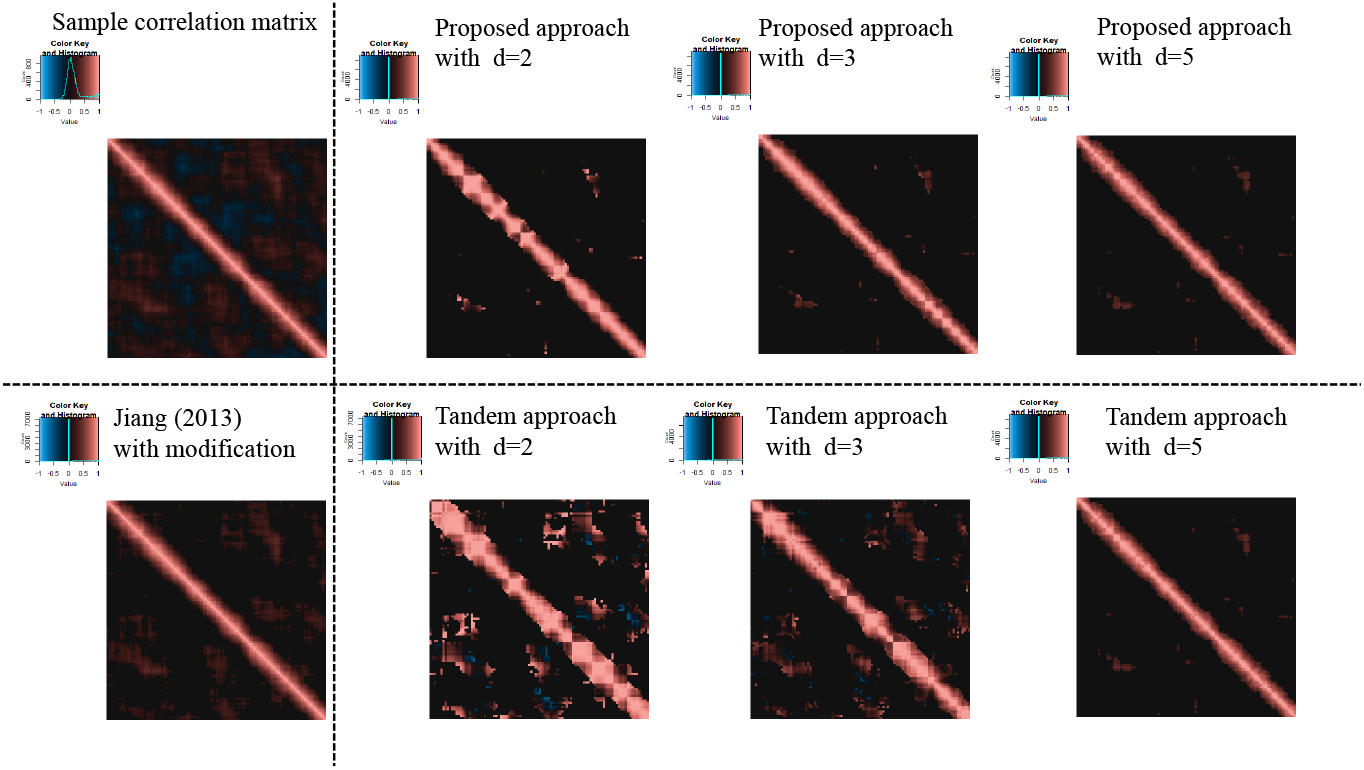
Examples of estimated correlation matrices for the true correlation model 1 (*n* = 75)

**Figure 14:**
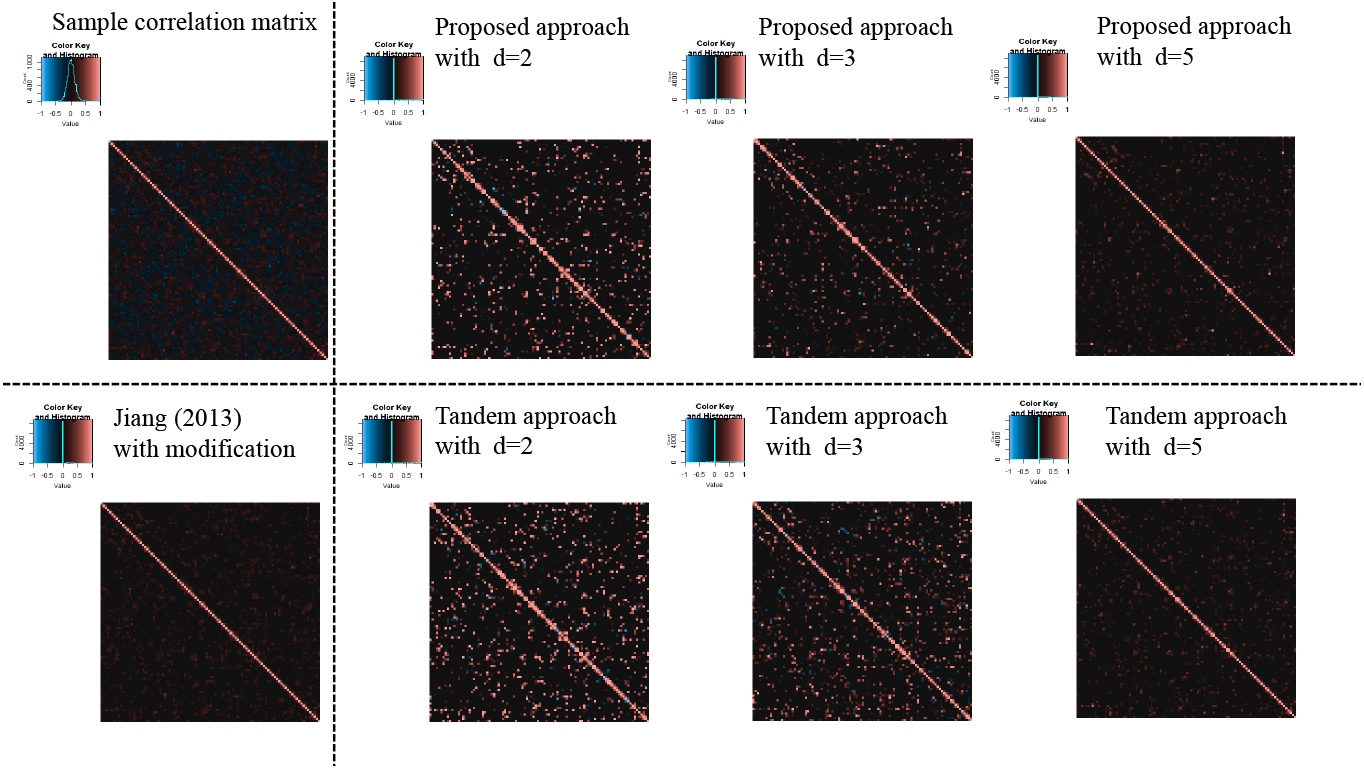
Examples of estimated correlation matrices for the true correlation model 2 (*n* = 75)

**Figure 15:**
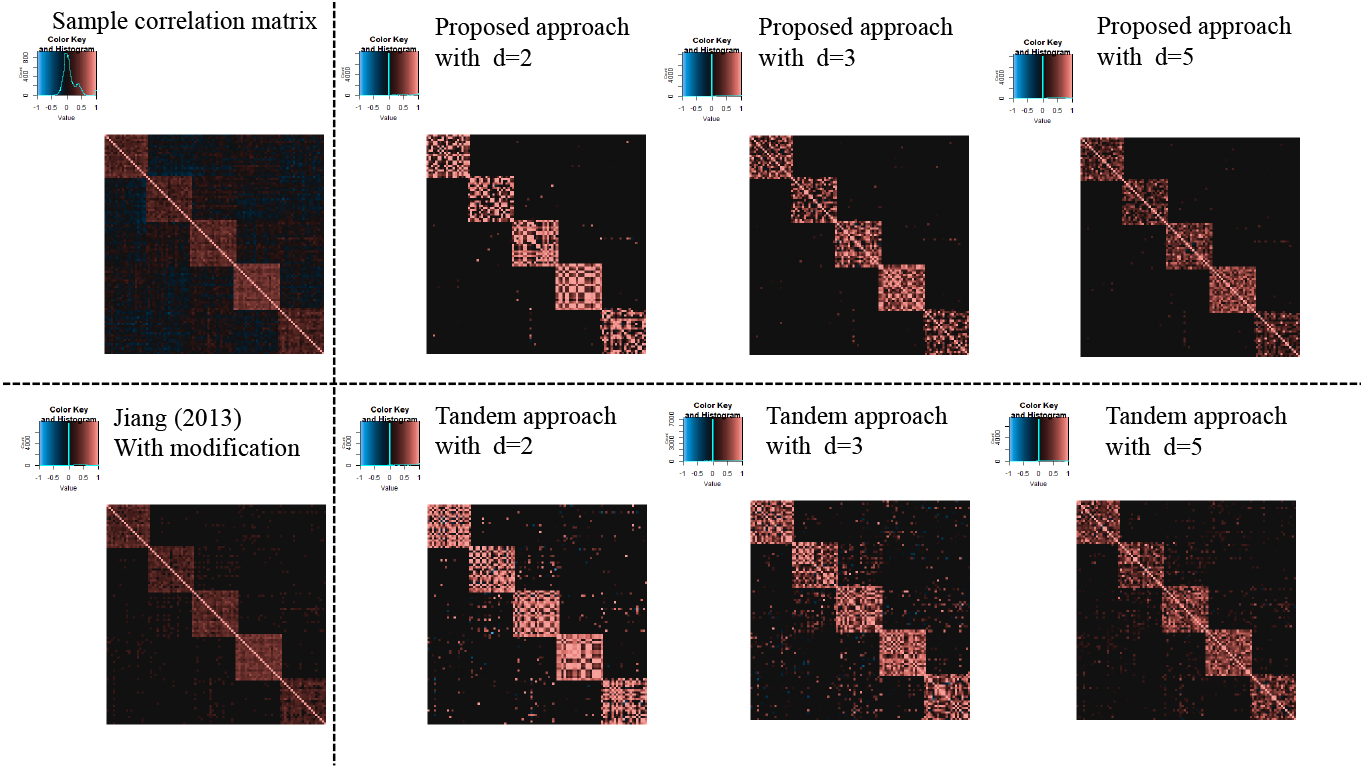
Examples of estimated correlation matrices for true correlation model 3 (*n* = 75)

### 3.2 Result of application of microarray gene expression dataset

In this subsection, the results of the application of the microarray gene expression dataset are shown. For the estimated original correlation matrix, Jiang (2013) with modification, the proposed approach and tandem approach, see Figure 16. Hence, the percentage points of *d* = 2, 3, and 5 in the proposed approach were estimated as *α* = 0.82, 0.81, and 0.75, respectively, while the percentage points in the tandem approach and Jiang (2013) with modification were both *α* = 0.65. The estimated results of Jiang (2013) with modification are as presented in Figure 16. However, FPRs were higher than those of the proposed approach. Here, the FPR is unaffected by the choice of rank in the tandem approach. From these results, the estimated sparse low-rank correlation matrix tends to be sparser when the rank is set as lower. In fact, it can be confirm that in Figure 16. In addition, as the rank is set larger, the estimated correlations of the proposed approach become like those of the tandem approach. We also confirmed that the estimated sparse low-rank correlation matrix between genes belonging to “informative” class tend to be similar to the results obtained in Rothman et al. [2009] using the heatmap.

**Figure 16:**
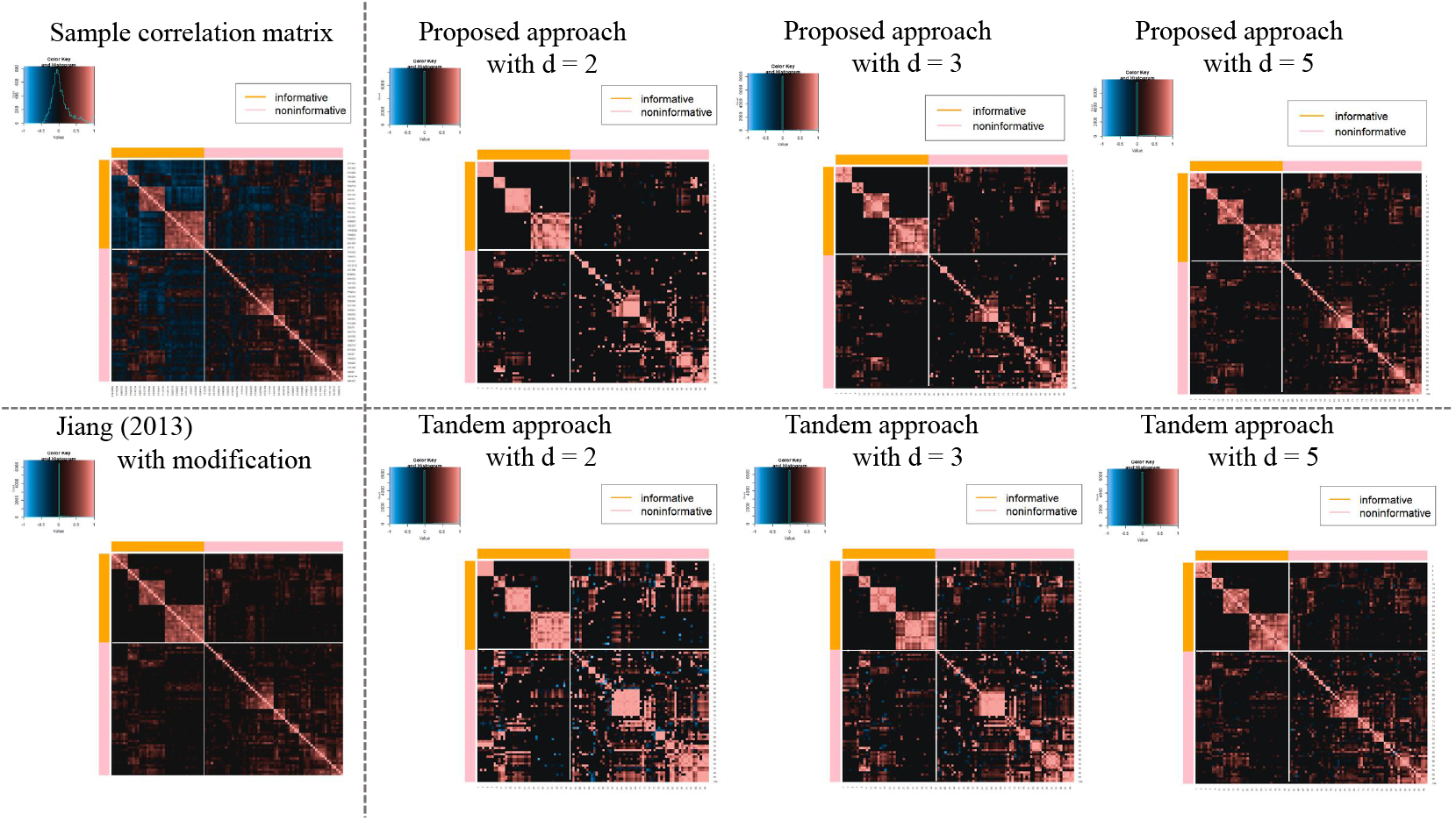
Estimated sparse low-rank correlation matrices with *d* = 2, 3 and 5, sample correlation matrix and sparse correlation matrix without rank reduction.

Next, Table 5 shows the results of the FPR of the proposed approach, tandem approach, and Jiang (2013) with modifications. The FPRs of the proposed method with *d* = 2, 3, and 5 were all lower than those of both the tandem approach and Jiang (2013) with modifications. In addition, the tendency can be confirmed from Figure 16 visually. In fact, we can confirmed that the proposed method was able to estimate the correlation coefficient between the classes as zero, compared to the tandem approach. The tendency was observed regarding the results of the numerical simulations.

**Table 5:** FPR of application of microarray gene expression dataset

| Method | $d$ | FPR |
| --- | --- | --- |
| Proposal | 2 | **0.128** |
| Proposal | 3 | 0.139 |
| Proposal | 5 | 0.197 |
| Tandem | — | 0.285 |
| Jiang (2013) with modification | — | 0.285 |

## 4 Conclusion

This study proposed a novel estimation method for sparse low-rank correlations based on the MM algorithm. The approach overcomes the problem of estimating low-rank correlation matrices. Low-rank approximation is an immensely powerful tool, and the approach provides us with an easy interpretation of the feature because the contrast of the estimated coefficients becomes larger. However, these estimation sometimes leads to misinterpretations. Simply put, even if the true correlation coefficient is zero, the corresponding estimated coefficient of the low-rank approximation without sparse estimation may be greater than zero. To confirm the efficiency of the proposed method, we performed a numerical simulation and conducted an experiment using a real example, which involved the use of a microarray gene expression dataset. In the case of the real example, the FPR of the proposed approach with *d* = 2, 3, and 5 were found to be 0.128, 0.139, and 0.197, respectively, although those of the tandem approach and Jiang (2013) with modifications were found to be 0.285 and 0.285, respectively. We were therefore able to confirm that the FPRs of the proposed approach is the best, irrespective of the rank. Similarly, from the numerical simulation, we confirmed that the FPRs of the proposed approach is superior to those of the tandem approach and Jiang (2013) with modifications. In addition to that, we also found that these relative errors of the proposed approach were superior to those of the tandem approach through the numerical simulations. From the result, the proposed approach is considered as approximating true correlation matrix compared to the tandem approach.

Although the proposed approach showed promising results on both the experiments, there are several aspects that need further investigations. First, when the rank is set to be low, the TPRs of the proposed method were low as compared to those of the tandem approach. The relationship between the determination of the rank and the corresponding TPR was a trade-off. Therefore, a method for a more effective determination of the rank should be developed. Second, in this study, the interpretability of the proposed method was not quantitatively evaluated. Therefore, the interpretability of the proposed method must be evaluated. We provide several examples of results estimated using heatmaps. From the visualization results, we were able to confirm the effectiveness of the proposed method; the results showed the proposed method was able to emphasize the features and lower the FPRs. Finally, it must be noted that the proposed approach only focuses on the positive correlation matrix and does not consider the negative correlation matrix for the sake of simplicity of interpretation. However, the proposed approach can be extended for considering both positive and negative correlation coefficients.

